# A systematic study of repetitive transcranial magnetic stimulation to enhance working memory manipulation abilities

**DOI:** 10.1101/278655

**Authors:** L. Beynel, S.W. Davis, C.A. Crowell, S.A. Hilbig, W. Lim, D. Nguyen, A.V. Peterchev, B. Luber, S.H. Lisanby, R. Cabeza, L.G. Appelbaum

**Affiliations:** Department of Psychiatry and Behavioral Science, Duke University School of Medicine, Durham, NC; Department of Neurology, Duke University School of Medicine, Durham, NC; Center for Cognitive Neuroscience, Duke University, Durham, NC; Department of Biomedical Engineering, Duke University, Durham, NC; Department of Electrical and Computer Engineering, Duke University, Durham, NC; Department of Neurosurgery, Duke University School of Medicine, Durham, NC; National Institute of Mental Health, Bethesda, MD; Department of Psychology & Neuroscience, Duke University, Durham, NC

**Keywords:** Transcranial Magnetic Stimulation, Online rTMS, Dorsolateral Prefrontal Cortex, Working Memory, fMRI, Electric Field Modeling

## Abstract

A core element of human working memory (WM) is the ability to perform mental operations on information that is stored in a flexible, limited capacity buffer. Given the profound importance of such WM manipulation (WM-M) abilities, there is a concerted effort aimed at developing approaches to improve them. Past research has identified neural substrates of WM-M centered in the dorsolateral prefrontal cortex (DLPFC), thereby providing a plausible and accessible target for noninvasive neuromodulatory stimulation that can be used to alter cortical excitability and potentially lead to facilitation of WM-M. In the current study, 5Hz online repetitive transcranial magnetic stimulation (rTMS), applied over the left DLPFC, was used to test the hypothesis that active rTMS would lead to significant improvements in memory recall accuracy compared to sham stimulation, and that these effects would be most pronounced in the WM-M conditions with the highest cognitive demand (registered Clinical Trial: #NCT02767323). Participants performed a delayed response alphabetization task with three individually-titrated levels of difficulty during active and sham rTMS. Analyses revealed that active rTMS led to numerically greater accuracy relative to sham stimulation for the hardest condition; however, this effect did not survive Bonferroni correction over all task conditions. Despite the lack of robust, study-wise significant effects, when considered in isolation, the magnitude of behavioral improvement in the hardest condition was negatively correlated with parametric difficulty-related fMRI activity in the targeted brain region, suggesting that individuals with less activation benefit more from rTMS. The present findings therefore suggest evidence towards the hypothesis that active rTMS can enhance performance during difficult memory manipulation conditions; however, firm conclusions cannot be drawn given the lack of overall significant effects. These findings are discussed in the context of individualized targeting and other factors that might moderate rTMS effects.

## Introduction

Working memory (WM) refers to interconnected processes that enable the temporary storage and online processing of information (Baddeley, 1998). A fundamental distinction in WM processes exists between the storage of information within WM buffers, or *maintenance*, and the active processing and reorganization of this information within WM, or *manipulation* (D’Esposito et al., 1999). With a few exceptions (e.g., keeping a phone number in mind until it is dialed), most WM tasks in real life require both maintenance and manipulation (e.g., translating a shopping list into the shortest path through your grocery store). In general, manipulation is sensitive to factors that affect brain function, including aging (Kirova et al., 2015) and a variety of psychiatric (e.g., Horan et al., 2008) and neurological disorders (e.g., Belleville et al., 2003). Given the critical role of WM in daily life, there has been a concerted effort to implement approaches to improve this ability. One such approach is non-invasive brain stimulation, such as repetitive transcranial magnetic stimulation (rTMS). Under this approach, the high intensity magnetic field of rTMS induces brief currents in the brain that modify cortical excitability and have been shown to vary with stimulation frequency, with higher frequencies (≥5Hz) generally increasing cortical excitability (Pascual-Leone et al., 1994). Correspondingly, fMRI studies have shown that high frequency rTMS can boost blood-oxygenation-level-dependent (BOLD) activity associated with successful performance during a host of different cognitive tasks, including episodic memory tasks (Vidal-Pineiro et al., 2014) and WM tasks (Esslinger et al., 2014). Importantly, high frequency rTMS has been shown to improve performance during some types of WM tasks. For example, there is evidence that ‘online’ rTMS (i.e., applied during the task) delivered to parietal cortex during spatial (Hamidi et al., 2008) and verbal (Luber et al., 2007) WM tasks can quicken reaction times to retrieve maintained information. Despite this promising evidence of rTMS enhancement of WM maintenance abilities, no studies have used online rTMS to enhance the critical skills underlying WM manipulation.

To address this gap, the current study tested whether it is possible to enhance behavioral performance during WM manipulation with rTMS by comparing the effects of active 5 Hz rTMS and sensory-matched electrical sham stimulation applied over the left dorso-lateral prefrontal cortex (DLPFC) during a delayed recall alphabetization task (DRAT). The DLPFC was chosen as the target for rTMS given past evidence of its involvement in the manipulation of information in WM. For example, previous studies have shown greater activity within the DLPFC during the delay period of a WM manipulation task than during WM maintenance task (D’Esposito et al., 1999, for a review see Curtis and D’Esposito, 2003). Moreover, past studies have shown that patients with DLPFC damage, compared to patients with non-DLPFC lesions or healthy subjects, present deficits in visual and verbal manipulation, without affecting performance in maintenance tasks (Barbey et al., 2013). In consideration of the frequency of rTMS stimulation, past studies have shown that online rTMS could induce performance enhancement by entraining endogenous task-related oscillatory dynamics. When applied at alpha frequency, online rTMS has been shown to induce a boost in alpha-power band at the targeted region (Thut, et al. 2011) along with corresponding behavioral performance enhancement (Klimesch et al., 2003). Given the important role of theta oscillations in memory processes (Roux, & Ulhas, 2014), and based on our previous results demonstrating that 5Hz rTMS resulted in performance enhancement to WM maintenance (Luber et al., 2007), 5 Hz was selected as the stimulation frequency in the present study. In order to further optimize the effectiveness of rTMS, individualized fMRI statistical parametric maps and electric field (E-field) models were used to define the stimulation target. Based upon this overall design, improvements in WM-manipulation (WM-M) abilities induced by active rTMS, over sham, were expected, especially in the most difficult trials. Indeed, some studies have suggested that rTMS effects occur only when the neural processes are challenged. For example, rTMS improvement has been found only for larger set sizes in a maintenance task (Luber et al., 2007; 2008; 2013), and only for degraded spatially filtered images and not images presented in their original resolutions during an object identification task (Viggiano et al., 2008). These goals were pre-registered under Clinical Trial number NCT02767323.

In addition to the primary goal of testing rTMS effects on WM-M, a secondary goal of this study was to evaluate factors that could moderate potential rTMS effect. As such, follow-up exploratory analyses were tested in order to evaluate the relationship between effect sizes, task-induced fMRI brain activations and rTMS stimulation dose. Regarding task-induced fMRI activity, it has been proposed that the initial activation state of the brain could influence TMS effect (Silvanto et al., 2008). However, there are two competing hypotheses in the literature regarding which brain activation state facilitates the rTMS effect. Using psychophysical adaptation paradigms to decrease the initial activation of neural populations, it has been shown that TMS behaviorally facilitates the more deactivated neural populations (Cattaneo and & Silvanto, 2008; Silvanto et al., 2007). On the other hand, some studies have found rTMS to be more effective when activity in the targeted region is high. For example, BOLD activity in the left premotor cortex increased when rTMS was applied to this area during voluntary hand contraction, though it decreased during rest (Bestmann et al., 2007). The same pattern has been replicated using somatosensory input (Blackenburg et al., 2008). Although these studies were based on sensory tasks, it is possible that brain state could also have an impact during higher level cognitive tasks such as the one employed here. As such, rTMS effect in the current study could correlate with parametric brain activations associated with increasing WM demand, either positively according to the results from TMS-fMRI studies or negatively according to Silvanto’s state-dependency results. Lastly, in order to evaluate the novel approach of targeting based on individual fMRI activations and E-field used in this study, additional analyses were performed to compare the Euclidian distance between individual and averaged coil location to determine if this is a predictor of rTMS effects.

## Methods

### Participants

Forty-nine healthy young adults were recruited and provided written informed consent for the study, which was approved by Duke University Institutional Review Board (IRB protocol #Pro00065334) and registered on ClinicalTrials.gov (NCT02767323). Across the 49 recruited participants, 5 of them were excluded during the screening session due to poor behavioral performance (n=3), or contra-indications to rTMS (n=2). Over the 44 remaining participants, 15 dropped out for scheduling reasons (n=6), noisy fMRI induced by movement (n=4), or due to pain induced by rTMS (n=5). Twenty-nine participants completed the full protocol. These 29 individuals had a mean age of 22.9 ± 4.8 years (from 18 to 35 years old) and consisted of 17 females and 12 males. Participants had normal, or corrected-to-normal, vision and were native English speakers. Participants were compensated $20/hour for their efforts with a $100 completion bonus.

### Experimental Protocol

The participants were scheduled for a total of 6 sessions. The first visit consisted of consenting, exclusionary screening, measurement of the resting motor threshold and 6 blocks of practice with the behavioral task. Participants then returned for an MRI visit and four more rTMS sessions (**Figure 1**). Visits 1, 2 and 3 were separated by approximately 1 week each and on average, participants completed the four rTMS visits in 11 days, on average.

**Figure 1:**
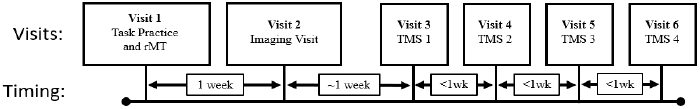
Schematic illustration of the full experimental protocol, describing the different visits and the relative time interval between each of them.

The first visit started with consenting and exclusionary screening wherein potential participants were excluded if they had current or past Axis I psychiatric disorders including substance abuse/dependences, as determined by the MINI International Neuropsychiatric Interview, English Version 5.0.0 DSM-IV (Sheehan and Lecrubier, 2002), or neurological disease as determined by the TMS Adult Safety Screen (Keel et al., 2001). All participants were also screened for substance use with urine drug screens and women of childbearing potential were screened with urine pregnancy tests. Individuals were excluded if they tested positive in either urine screen.

### Delayed-Response Alphabetization Task

The delayed-response alphabetization task (DRAT: **Figure 2A**) investigated WM manipulation processes by having participants reorder letters presented in a random sequence by alphabetizing them within WM. In this task, an array containing 3 to 9 letters was presented on a screen for 3 seconds, followed by a 5-second delay period during which the subjects were asked to keep this array in mind (maintenance) and to reorganize the letters into alphabetical order (manipulation). The set of letters included all English consonants, with vowels excluded to reduce spontaneous chunking. After the delay period, a letter with a number above it appeared on the screen for 4 seconds and participants were asked to report, using one of three button options, if the probe letter was not in the original set (New), if the letter was in the original set and the number matched the letter serial position when the letter sequence was alphabetized (Valid), or if the letter was in the original set but the number did not match the letter serial position when alphabetized (Invalid). Distinguishing between Valid and Invalid required successful alphabetization. The three conditions occurred in a random order with 40% of trials in the Valid condition, 40% in the Invalid condition, and 20% in the New condition. For all three conditions, the probe was never from the first half of the alphabetized array, and in the Invalid condition, to exclude obvious differences between correct and incorrect position, the number above the letter was always within 1 step of the letter’s actual alphabetized position. The response phase was followed by a 5-second inter-trial interval during which the participants got feedback displayed as “Correct” or “Incorrect”, in green or red font respectively, depending on their performance during the previous trial during practice trials.

**Figure 2:**
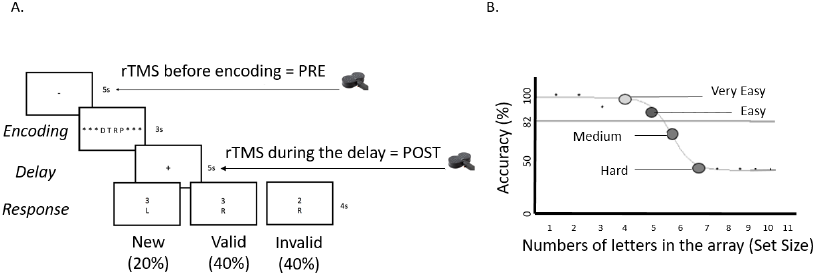
A) Schematic illustration of DRAT. One trial is shown with an array of 4 letters to encode, followed by a 5s delay period, during which subjects had to maintain and reorganize the letters into alphabetical order. Examples of the 3 possible responses are shown at the bottom: “New”: the letter was not in the original array; “Valid”: the letter was in the array and the number represented the correct position in the alphabetical order; “Invalid”: the letter was in the array but the number did not match the correct serial position when alphabetized. B) A schematic example of titrated task data and a fitted sigmoidal function used to define the individual difficulty levels with Very Easy and Easy defined as the set sizes for which the accuracy was greater than 82% and Medium and Hard as accuracy lower than 82%.

In their first visit, participants performed 6 blocks (150 trials) of the DRAT in order to estimate easy to hard set size levels for each individual. Twenty-five trials were included in each block with a brief, self-paced rest interval between blocks. During this first visit, the DRAT was performed using a 2-down-1 up staircase procedure: for each correct response the set size in the array was increased by 1, while it was decreased by 2 for each incorrect response. In order to define individual task difficulty levels for each participant, performance accuracy data (% correct) from the Valid and Invalid conditions of Visit 1 was fitted to a sigmoid function and subsequent set sizes were defined relative to 82% correct, the theoretical convergence point for a 2-down-1-up staircase. To assure that the psychometric function was not strongly influenced by low numbers of trials at the easiest and hardest set sizes, 50% accuracy was used for the largest set sizes if less than 10 trials were tested. To achieve more stable curve fits, anchor points were added to both ends of the set size by accuracy plots, using set sizes of 1 and 2 at 100% accuracy and set sizes 10 and 11 at 50% accuracy. The intersection between the 82% accuracy threshold and the fitted sigmoidal function was then identified as the breakpoint for subsequent set size assignments. The two set sizes lower than the intersection values were defined as the Very Easy and Easy levels, while the two set sizes greater than this value were defined as Medium and Hard (**Figure 2B**). All four individualized levels were used in the subsequent MRI session, but the Very Easy level was not included in the TMS sessions to avoid ceiling effects and increase the number of trials per condition in the study.

### MRI Acquisition

Participants were scanned on a 3-T gradient-echo scanner (General Electric 3.0 Tesla Signa Excite HD short bore scanner), equipped with an 8-channel head coil. During this session, a structural MRI and a diffusion weighted imaging (DWI) scan were acquired, as well as functional acquisitions while subjects performed 4 blocks of the DRAT. The anatomical MRI was acquired using a 3D T1-weighted echo-planar sequence (matrix = 256^2^, time repetition [TR] = 12 ms, time echo [TE] = 5 ms, field of view [FOV] = 24 cm, slices = 68, slice thickness = 1.9 mm, sections = 248). 3D T2-weighted, with fat saturation, echo planar sequence were also acquired (matrix = 256^2^, TR = 4000 ms, TE = 77.23 ms, FOV = 24 cm, slice thickness = 2 mm). Coplanar functional images were acquired using an inverse spiral sequence (64 × 64 matrix, TR = 2000 ms, TE = 31 ms, FOV = 240 mm, 37 slices, 3.8-mm slice thickness, 254 images). Finally, DWI data were collected using a single-shot echo-planar imaging sequence (TR = 1700 ms, slices = 50, thickness = 2.0 mm, FOV = 256 × 256 mm^2^, matrix size 128 × 128, voxel size = 2 mm^3^, b value = 1000 s/mm^2^, diffusion-sensitizing directions = 25, total images = 960, total scan time = 5 min).

Stimuli for the DRAT were back-projected onto a screen located at the foot of the MRI bed using an LCD projector. Subjects viewed the screen via a mirror system located in the head coil and the start of each run was electronically synchronized with the MRI acquisition computer. The DRAT was performed using the 4 titrated difficulty levels defined from Visit 1. Overall accuracy was presented on the screen at the end of each block of 30 trials. Behavioral responses were recorded with a 4-key fiber-optic response box (Resonance Technology, Inc.). Scanner noise was reduced with ear plugs, and head motion was minimized with foam pads. When necessary, vision was corrected using MRI-compatible lenses that matched the distance prescription used by the participant. The total scan time, including breaks and structural scans, was approximately 1 hour 40 minutes.

### MRI Processing

Functional images were preprocessed using FSL image processing tools, including FLIRT and FEAT, in a publically available pipeline developed by the Duke Brain Imaging and Analysis Center (https://wiki.biac.duke.edu/ biac:analysis:resting_pipeline). Images were skull stripped, reoriented and corrected for slice acquisition timing, motion, and linear trend; motion correction was performed using FSL’s MCFLIRT, and 6 motion parameters were then regressed out of each functional voxel using standard linear regression. Images were then temporally smoothed with a high-pass filter using a 190s cutoff, and normalized to the Montreal Neurological Institute (MNI) stereotaxic space. Separate events were modeled for the array presentation (duration: 3s), delay period (duration: 5s), and response (duration: subject response time), each with an onset at the beginning of the event. Incorrect and non-response trials were modeled identically, but separately.

Parametric statistics were used to examine changes in the neural correlates of underlying WM manipulation, allowing activity to be modeled as a function of discrete changes associated with set size increases. These adaptive changes allowed us to model how responsive an individual would be to parametric variability in the set size across trials. At the first level, functional data were analyzed as individual runs. Second-level analyses combined data across runs for each subject using a fixed-effects model. Functional data were analyzed using a general linear model (GLM) in which trial events were convolved with a double-gamma hemodynamic response function. The GLM examined BOLD response during trials where the correct response was chosen in the behavioral task. The GLM included separate regressors modeling the duration of the array, the duration of the delay period, and the duration of the probe period until the time of the subject’s response. Additionally, weighted regressors were included during the delay period to model the parametric increase in difficulty with increased set size. These were orthogonalized with the delay period regressor. This processing allowed for the identification of individualized statistical maps that predicted the parametric increase in BOLD activity associated with increasing set size. The peak of activation within the left medial frontal gyrus in each subject was chosen as the rTMS target and entered into the neuronavigation system (BrainSight, Rogue Research, Canada).

### Electric field modeling and TMS targeting

The effects of TMS coil positioning depend on the individual head anatomy and the spatial distribution of the induced E-field, which determines what neural populations are affected (Peterchev et al., 2012). Thus, simulation of the E-field induced by TMS in individual subjects is increasingly recognized as an important step in spatial targeting of specific brain regions, and forms a critical link between the externally applied rTMS parameters and the neurophysiological response that supports cognitive operations. To determine the E-field induced in the brain by TMS for each subject, simulations using T1, T2, and DWI images and the finite element method in the SimNIBS software package (Windhoff et al., 2013) were conducted. The models featured five distinct tissue types: skin, skull, cerebrospinal fluid, grey matter, and white matter. The DWI information was used to generate anisotropic conductivities for white matter using the volume-normalized approach. The spacing between the coil and the scalp was assumed to be 4 mm (the default for SimNIBS).

To select the position of the figure 8 coil for rTMS across 3 locations and 3 angular parameters, E-field models were used to determine the maximum overlap between the E-field strength and the activations from the fMRI task (see **Figure 3**). The optimization focused on the E-field strength since it appears to be the key determinant of the neural recruitment by TMS (Bungert et al., 2017). The E-field was simulated at 54 coil targets (9 positions and 6 orientations per position) with a model of the figure 8 coil used in the study (A/P B65 Coil, MagVenture, Denmark). The 9 positions were generated by placing a 3 × 3 grid with 1 cm^2^ spacing above the peak fMRI activation. For each position, 6 different coil orientations were simulated corresponding to 30° rotation increments in a 180° semicircle. Due to the symmetry of the E-field, the 180° semicircle was sufficient to encompass all orientations in the full 360° circle. The E-field magnitude distributions, constrained with a magnitude higher than 0.4 V/m, for each of the 54 coil targets were correlated with the z-values of the fMRI activation. The coil position and orientation with the highest correlation was selected as the primary rTMS target, while the second and third highest correlations were noted as backup targets in the event that the primary target could not be used, for instance due to tolerability issues.

**Figure 3:**
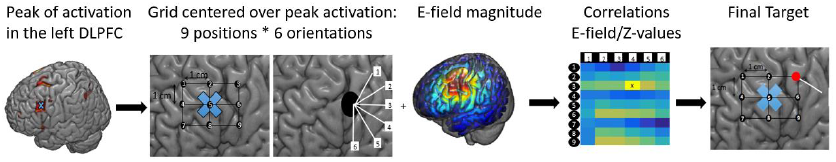
TMS targeting procedure illustrated for one subject. From left to right: Peak BOLD activation (1<z<3) on the left DLPFC associated with increasing set size; E-field grid with 3 × 3 positions and 6 coil orientations centered at the peak BOLD activation; representation of the E-field magnitude for one of the 54 options; correlation matrix between E-field magnitude for each of the 54 options and the activation z-values in which the highest correlation is for position 3 and orientation 4; optimized rTMS coil location (red dot) and orientation (white line).

### TMS procedures

#### Motor threshold determination

After the initial screening visit, hot spot determination for the right first dorsal interosseus (FDI) representation in motor cortex and resting motor threshold (rMT) were assessed for each participant. TMS was performed with an active/placebo figure-of-8 coil (A/P B65) and a MagPro X 100 stimulator with MagOption (MagVenture, Denmark), while the coil position was continually monitored through a stereotaxic neuronavigation system (Brainsight, Rogue Research, Canada). The TMS device was configured for biphasic pulses, in standard pulse width mode with the direction of the TMS pulses such that the initial phase of the E-field pointed from anterior to posterior direction. Electrodes (Neuroline 720, Ambu) were placed on the FDI in a belly–tendon montage and motor evoked potentials (MEPs) were recorded by an electromyogram (Power Lab and LabChart). The motor hot spot was defined as the position over the left motor cortex that elicited the greatest MEP in the right FDI. The resting motor threshold (rMT) was then defined as the TMS pulse intensity producing on average an MEP of 50 μV peak–peak amplitude, using a maximum likelihood method (TMS Motor Threshold Assessment Tool, MTAT 2.0, Awiszus, 2003).

#### Repetitive TMS

Subsequent to defining a stimulation target based on data from fMRI activations and E-field modeling, described above, participants returned for visits 3 through 6 in which they performed the DRAT while active or sham rTMS was delivered to the left DLPFC target. Twenty-five pulses of 5 Hz rTMS were delivered at 100% of rMT on each trial, either immediately before encoding (Pre) or post-encoding, during the delay period of the DRAT (Post), see **Figure 2A**. Sham stimulation was applied using the same coil (A/P B65) in placebo mode, which produced similar clicking sounds and somatosensory sensation (via electrical stimulation with scalp electrodes) as in the active mode, but without a significant magnetic field reaching the brain. Coil position was set according to the E-field optimized dosing using the BrainSight neuronavigation system and was maintained at a high level of precision throughout the session with real-time robotic guidance using the Smart Move Robot (Advanced Neuro Technology, Netherlands). On each visit, subjects performed the DRAT with the titrated 3 difficulty levels (Easy, Medium, and Hard), with feedback on the overall accuracy given at the end of each block. Ten blocks of the DRAT task were performed: a first block without stimulation (NoStim1), four blocks of active or sham stimulation, one block without stimulation (NoStim2), and four more blocks with the sham or active stimulation. The order of stimulation type (Active or Sham) was presented according to an ABBA schedule that repeated twice (one cycle during the first two sessions and one cycle during sessions three and four), while the stimulation timing (Pre or Post) alternated from block-to-block and flipped order between the 2^nd^ and 3^rd^ session. The starting order for stimulation type and stimulation timing was counterbalanced across participants.

#### Analysis of Behavioral, Neural, and Stimulation Dose Determinants of rTMS Effects

The main analyses of interest in this study concerned the influence of rTMS on WM manipulation abilities. In order to infer these relationships, several analytical steps were taken. First, behavioral performance was defined by focusing on trials and conditions of interest. As such, data from the first three trials of each block were removed, as participants were asked to provide feedback on the somatosensory effects of rTMS and thus did not focus on the task. Although the identity of a trial as Valid, Invalid or New is only determined by the probe and even though subjects have to perform the same alphabetization for all conditions, data from all ‘New’ probe trials were removed as they had been included as catch trials. Furthermore, given that the difficulty levels were individually titrated, difficulty level was normalized according to the starting set size for each participant. The reaction times were not emphasized during this experiment, and reflect more the decision process than the WM manipulation; the analyses were therefore performed on accuracy only. Using behavioral accuracy data from the remaining trials, preliminary analysis was performed to test if any cumulative within-session effects of rTMS occurred. To do so, ANOVA was conducted comparing accuracy before any stimulation to accuracy after active rTMS and after sham rTMS. Next, to assess rTMS effects on accuracy during the DRAT task, 2 × 4 × 3 × 2 × 2 repeated measures ANOVA was performed with the following within-subject factors: *Condition* (Valid vs. Invalid), *Visit* (1, 2, 3, and 4), *Difficulty* (Easy, Medium and Hard), *Stimulation Type* (Active and Sham) and *Stimulation Timing* (Pre and Post). When appropriate, post-hoc comparisons were performed using Bonferonni correction.

A second analysis was conducted to explore the relationships between brain activations, stimulation intensity, E-field magnitude, and rTMS effects. To define brain activations for the targeted ROI, subject-specific MNI-space brains were created by an affine registration between the MNI T1 2mm brain template using FSL’s FLIRT. The MNI subject-specific brains then underwent another affine registration to the Harvard-Oxford 471 ROI templates. The z-values obtained from the parametric increase in BOLD activity associated with increasing set size during the delay period were then extracted for each of these ROIs. Finally, the z-values from the targeted ROI were defined according to the correspondence between target coordinates registered on BrainSight and ROI number defined by MRIcron.

To examine whether E-field exposure affected rTMS outcomes, the center of mass (CoM) of the E-field for the voxels with positive z-values within the left DLPFC was calculated as follows:

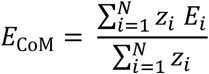

where *N* is the total number of voxels within the left DLPFC with positive z-values, and *z*_*i*_ and *E*_*i*_ are the z-value and E-field magnitude for voxel *i*, respectively. This definition assigns higher weights to E-field values corresponding to stronger fMRI activations, to compute a composite measure that is most relevant to the fMRI activations.

The obtained z-values and E-field measures were then correlated with the rTMS effect, calculated as follows:

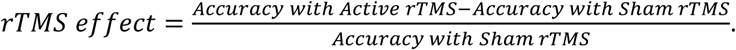

## Results

### Tolerability

In addition to the five subjects who stopped participation in the study because of pain, deviations from the planned stimulation protocol occurred for nine other subjects who completed the full study. For seven of these participants, the intensity of rTMS was reduced to alleviate discomfort (with relative intensities of 82, 88, 95, 96, 98, 98 and 98% of the initial measured rMT). In addition, for 4 subjects (2 of whom also required a reduced intensity level), the placement of the coil was adjusted by selecting the backup e-field target (n=2), the second backup E-field target (n=1) or the target defined by fMRI activations rather than the target defined by the E-field (n=1). Because performance of these subjects were not outliers from group means, they were kept in the analysis. However, to take these deviations into account, subsequent analyses were performed on the adjusted E-field exposure and stimulation intensity rather than the initial values derived from resting motor threshold.

### Cumulative effect of rTMS

The current design consisted of both active and sham stimulation, as well as two blocks of trials in which no stimulation was delivered. As a first step, analyses were performed to evaluate if active or sham stimulation produced a cumulative “carryover effect” on the blocks of trials with no stimulation. For this purpose, accuracy was collapsed across Visits and Difficulty Levels as a cumulative effect of rTMS was not expected to vary across these conditions. No cumulative effect of rTMS exposure (active or sham) was found in the accuracy data for the noStim blocks. ANOVA did not reveal any change in accuracy between blocks performed before stimulations (NoStim1: 73.14 ± 8.23%), blocks performed after active rTMS (NoStim2_afterActive: 71.18 ± 8.41 %), or blocks performed after sham rTMS (NoStim2_afterSham: 71.39 ± 8.39 %; F(2, 56)< 1). This result suggests that no carryover effects persisted following the blocks of trials in which active or sham stimulation was delivered.

### rTMS effects

To assess rTMS effects on accuracy during the DRAT task, an analysis was performed on performance obtained during the blocks with stimulation to determine whether active rTMS could enhance working memory manipulation abilities over sham rTMS. A first ANOVA including Stimulation Timing (Pre vs. Post) did not yield a significant main effect (F(1,27)<1) or interaction (F(1,27)<1) involving this factor, and hence, we collapsed across the Pre and Post conditions. The resulting two (Condition: Valid and Invalid) by four (Visit: 1, 2, 3, 4) by three (Difficulty: Easy, Medium, Hard) by two (Stimulation Type: Active, Sham) repeated-measures ANOVA revealed a main effect of Condition (F(1,28)=7.9, p<.01), showing that subjects performed significantly better in the Valid condition (73.28 ± 8.2 %) than in the Invalid condition (63.81 ± 8.9 %). The ANOVA also revealed significant main effects of Visit (F(3, 84)=9.87, p<.01) and Difficulty (F(2,56)= 177.01, p<.01) (**Figure 4**). Post-hoc analyses comparing accuracy during the different Visits revealed that accuracy in the third (73.4 ± 8.2) and fourth visits (73.3 ± 8.2 %) were higher than in the first (69.1 ± 8.5 %) and the second (70.3 ± 8.4 %) (all ps < .01). Post-hoc analyses comparing accuracy over the difficulty levels revealed significant reductions in accuracy with larger set size for all pair-wise comparisons (p<.01).

**Figure 4:**
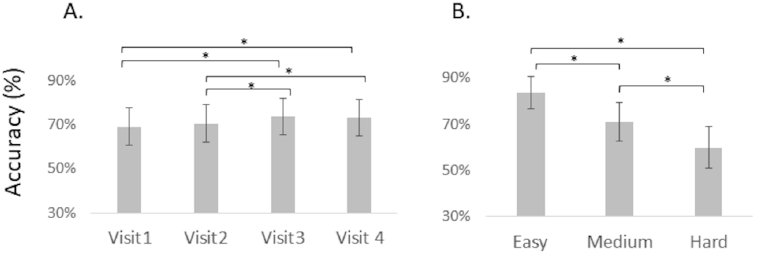
A) Accuracy across Visits. B) Accuracy across Difficulty Levels. Error bars represent between-subject standard errors, while statistics were performed as within-subject contrasts.

The ANOVA did not reveal a significant main effect of Stimulation Type (F(1,28)<1), and no significant two-way interaction between Difficulty and Stimulation Type (F(2, 56)=2.25, p=.12). However, there was a significant three-way interaction between Condition, Difficulty, and Stimulation Type (F(2, 56)=4.13, p=.02). To clarify this interaction, we conducted separate four (Visit) by three (Difficulty) by two (Stimulation Type) ANOVAs for the Valid and Invalid conditions. Results showed a significant interaction between Difficulty and Stimulation Type for the Invalid condition (F(2,56)= 5.15, p <.01), but not for the Valid condition (F(2,56)<1). Thus, consistent with our expectations, rTMS effects were more pronounced in the most difficult trials.

Post-hoc analyses (**Figure 5A.**) revealed that subjects were numerically more accurate with active than with sham rTMS at the hardest difficulty level (52.3 ± 9.3 % vs. 48.3 ± 9.3 %) though this did not reach statistical significance at a Bonferroni corrected alpha level of .05 (F(1,28)=11.21, p=.12). In contrast, performance was comparable for the two stimulation conditions for the easy (77.3 ± 7.78 % vs. 79.77 ±7.5 %) and medium difficulty levels (62.2 ± 9.0 % vs. 62.8 ± %) (F(1,28)=1.42, p =1; F(1,28)< 1). To further investigate rTMS effects in the Invalid condition, active minus sham differences for each individual were tested using binomial statistics. For the hardest difficulty level, 21 out of 29 subjects showed accuracy improvement with active rTMS relative to sham (p=.024), while the probability levels did not reach significance in the easy or the medium difficulty levels, where both conditions had and 13 out of 29 subjects showing better accuracy with active rTMS (p= .71) (**Figure 5B**). The finding that rTMS was more effective at higher levels of difficulty is consistent with previous findings by Luber and collaborators (Luber et al., 2007; 2008; 2013) and confirm our a priori expectations.

**Figure 5:**
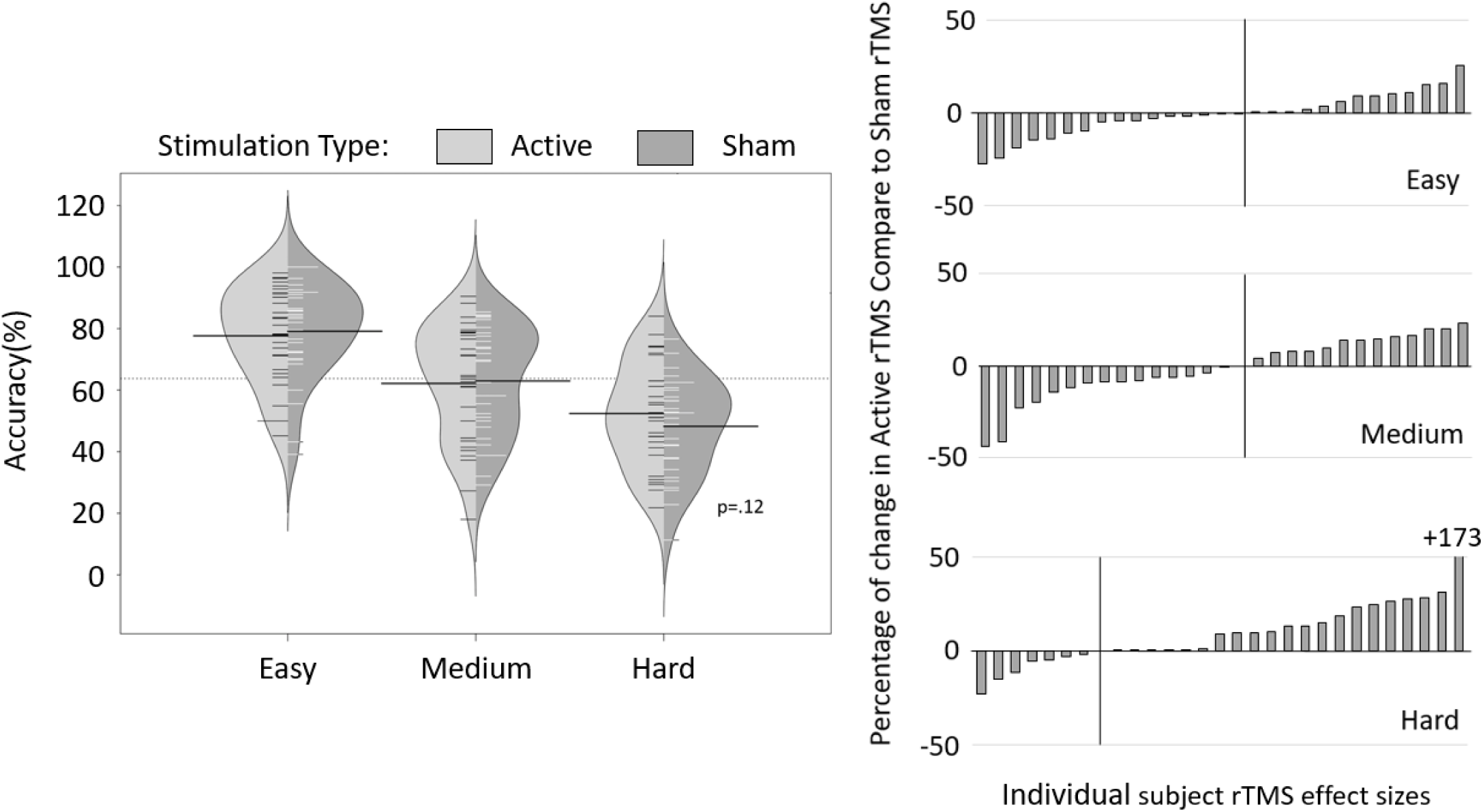
A) Mean Accuracy in the Invalid condition for active (light grey) and sham (dark grey) rTMS and for each difficulty Level. Small lines represent individual data and shaded curves the normal density trace. The darker lines represent the average for each condition and the light grey dashed line represent the overall average of accuracy B) Percentage of change in accuracy for active compare to sham rTMS for each individual subject for each difficulty level. Positive values indicate improved accuracy for active relative to sham stimulation.

### Exploratory analysis of potential moderators of rTMS effects

To assess if brain activation or stimulation dose could predict the magnitude of accuracy differences for active versus sham stimulation in the hardest difficulty level in the Invalid condition, where the performance slightly increased with active rTMS, an exploratory multiple regression was performed. For the *brain activations*, the z-values obtained from the parametric increase in BOLD activity associated with increasing set size, during the delay period were used. For the *stimulation dose, E*_CoM_, and intensity of stimulation were used. Although these two measures were highly correlated (r = 0.76, p<.001s), they relate to a different stimulation dose aspect: the dose within the presumably relevant region of left DLPFC, and the stimulation of nerves in the scalp, respectively. One subject showed performance improvements greater than 2 standard deviations above the mean and was excluded from this analysis. Analysis on the remaining 28 subjects revealed that the stimulation dose and brain activation indeed predicted the magnitude of accuracy differences for active versus sham stimulation, with the multiple regression model explaining 28% of the total variance (p<.04). Within this model, brain activations obtained from fMRI-derived parametric z-values were the only significant predictor of the accuracy difference (b = −5.33, CI= [-8.93; −1.75], b* = −0.64, t_24_= −3.07, p<.01). This significant relationship between activity and behavioral change indicates that lower z-values in the targeted ROI are associated with larger active-versus-sham differences (see **Figure 6**). This finding is consistent with state-dependency hypothesis that rTMS is more effective for the most deactivated neural populations (Silvanto et al., 2008), and extends available evidence for this to a higher cognitive task. The analysis did not reveal any relationship between *E*_CoM_ and accuracy differences (b = 1.09, CI = [-0.12; 2.03], b* = 0.6, t_24_= 1.84, p=.08), or between intensity of stimulation and accuracy differences (b= −0.83, CI = [-2.09; 0.42], b* = 0.41, t_24_= −1.38, p = .18).

**Figure 6:**
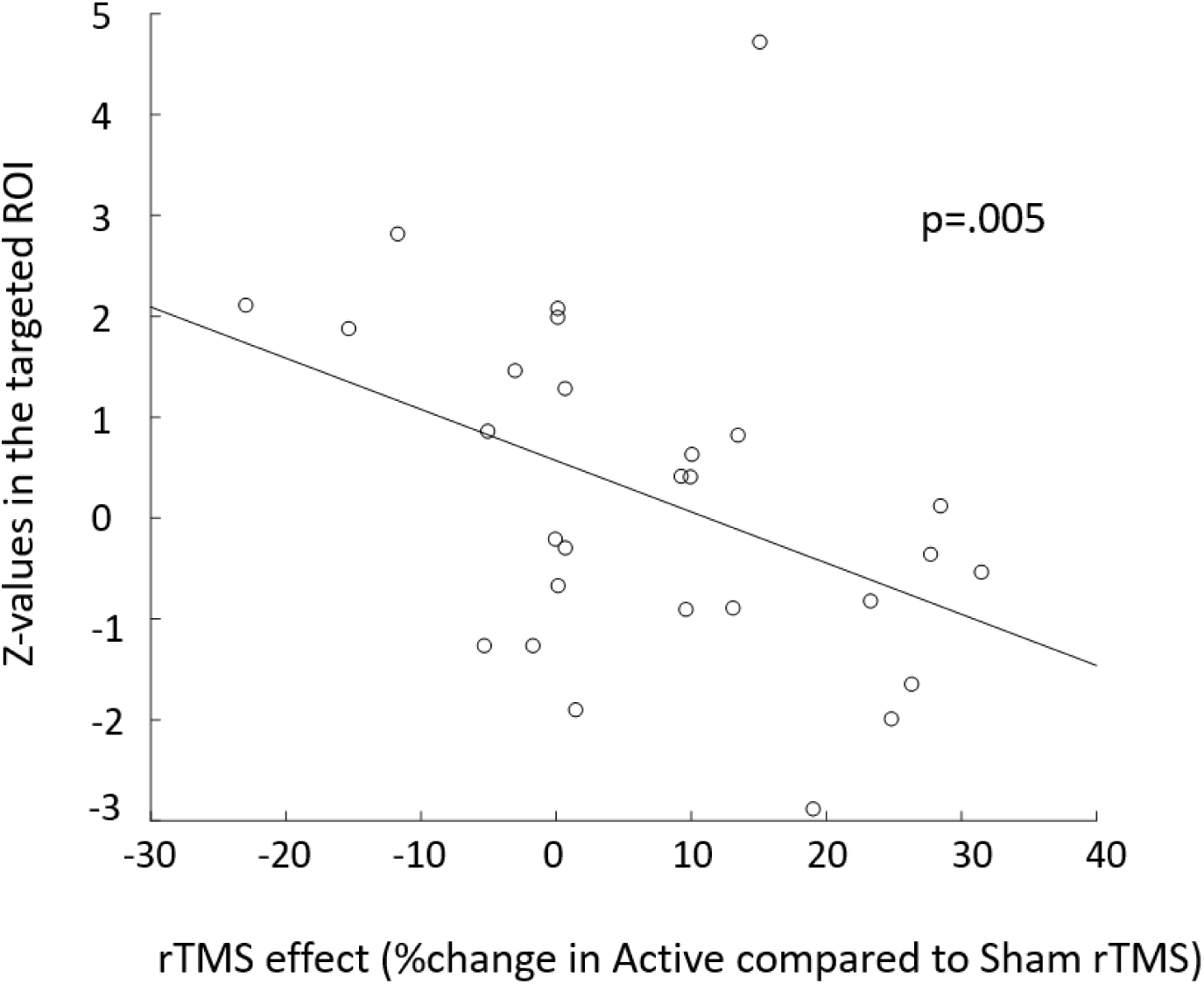
Scatterplot between z-values in the targeted ROI and difference between accuracy in the active versus sham conditions at the hardest difficulty level in the Invalid condition (b*=-0.64, t_24_= −3.07, p=.005).

To further investigate how individualized targeting could have impacted behavior, individual-subject target locations are depicted in Figure 7. Here target locations are shown on a standardized cortex and scaled according to the magnitude of active versus sham behavioral differences. As can be seen, the individualized fMRI activations combined with E-field modeling leads to a wide spatial distribution of stimulation targets. When considered in relation to the group mean target location (shown by the red sphere) which is centered in the MFG, several of the more remote subjects produce negative behavioral differences (sham > active). Therefore, in order to evaluate how the spread of individualized target locations impacts behavioral accuracy, the Euclidian distance between the group average and individual targets was calculated. This calculation was done for both the coil locations and the peak fMRI activations. No combination of these Euclidian calculations revealed significant correlations with behavioral differences for active versus sham stimulation (p > .20 for all comparisons), suggesting that the spatial spread of TMS targets does not explain the lack of significant rTMS effect in this study.

**Figure 7:**
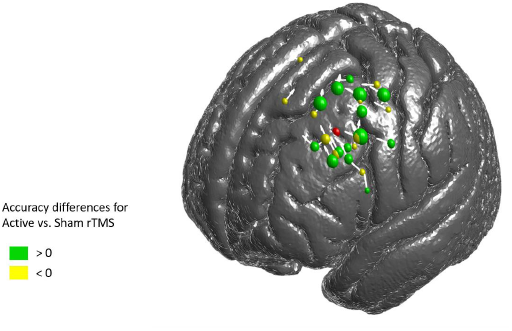
Coil position (sphere) and orientation (white arrows) for each subject. The color of the spheres represents the accuracy difference for active versus sham stimulation. The green spheres represent a stronger accuracy with active rTMS, while the yellow spheres represent a stronger accuracy with sham rTMS. In each case, the size of the sphere represents the magnitude of the rTMS effect. White arrows correspond to the direction of the first phase of the induced E-field pulse (some of the arrowheads are not visible because of the 3D view). The red sphere represents the average coil location across all subjects.

## Discussion

WM manipulation has been shown to be a critical cognitive capability that declines naturally with age and is profoundly impacted by neurodegenerative disorders and brain injury. Given the importance of this ability and past research showing promising effects of rTMS on cognition, the current study tested whether online rTMS could enhance WM manipulation abilities, which to our knowledge, has never been tested. Given the large involvement of the DLPFC (D’Esposito et al., 1999) and of theta (4-7 Hz) neural oscillations (Roux and Uhlhaas, 2014) in WM manipulation, 5Hz rTMS was applied over the left DLPFC in an attempt to determine if this stimulation was able to enhance performance on a difficulty-titrated WM manipulation task. Although the results did not survive Bonferoni correction, they revealed an expected pattern with slight improvement associated with active rTMS over the sham stimulation only in the hardest condition (i.e., the hardest difficulty level of the Invalid condition), where subjects performed significantly worse than in the other conditions. This pattern is in line with previous results that found rTMS improvement only in the most difficult conditions: the largest set sizes in a WM maintenance task (Luber et al., 2007; 2008; 2013), and blurred versus high-resolution images during an object identification task (Viggiano et al., 2008). Active rTMS was able to affect performance only at difficulty levels with the most room for improvement, indicating that elements of the task or the rTMS parameters could be optimized to further promote this benefit and produce globally significant study-wise effects. The following discussion addresses the strengths and weaknesses of this protocol while offering suggestions to improve future studies.

As a first consideration in an intervention study of this nature, it is important to discuss participant compliance and adherence to the planned protocol. In this study, a planned sample of 30 participants was sought to receive active rTMS at 100% of resting motor threshold and somatosensory matched electrical sham stimulation. To obtain these individuals, forty-four participants were enrolled (after passing screening and exclusionary criteria), 15 of these participants dropped out for scheduling reasons, noisy fMRI induced by movement, or due to pain induced by rTMS. In the end, this led to a final analysis sample of 29 participants, which is one less than the desired sample size but considerably larger than the typical sample size for online rTMS studies – mean of 13 across 143 studies of online TMS included in a current meta-analysis from our group (Beynel et al., in preparation). Despite this large sample size, sensitivity to detect significant effects may have been reduced due to the high number of conditions in the design (Visits [4], Difficulty [3], Stimulation Type [2], Stimulation Timing [2], and Conditions [2] after excluding catch trials from the ‘New’ condition). In the future, studies may wish to focus on a reduced the number of conditions to increase the effect size. Despite this, the difficulty levels used in the task were defined through a staircase procedure such that all subjects performed the task at difficulty levels adjusted according to their own behavioral performance. This approach allowed for a better between-subjects comparison and prevented the use of arbitrary set sizes where some subjects would perform at ceiling and others at chance.

With regard to the observed behavioral results, statistical consideration of participant accuracy revealed a main effect of *Condition*, with subjects performing significantly better in the Valid condition than in the Invalid condition. This result is in line with the cognitive cost associated with the rejecting familiar items, which requires inhibitory control (Jonides et al., 1998; Jonides & Nee, 2006). A main effect of *Visits* was found suggesting that participants learned to perform better with more experience at the task. This effect did not interact with Stimulation Type, implying that the stimulation parameters used in this study did not differentially affect learning. A main effect of *Difficulty* was also found, confirming that the staircase procedure used to define set sizes for each subject was appropriate and that subjects performed worse with larger set sizes. The results did not reveal any main effect of *Stimulation Timing* indicating that stimulation before the encoding period and during the delay period did not differentially affect accuracy. This null finding is consistent with results from EEG recording during a delayed match-to-sample task (Raghavachari et al., 2001) which revealed that theta oscillations increased dramatically at the beginning of the trial, continued through the delay period, and decreased at the end of the trial. The stability of the theta oscillations before and during the trial could explain why, in the current study, no differences were observed between stimulation applied at the two intervals. Another possibility is that participants started to rearrange the letters into alphabetical order during the array presentation. Finally, results failed to reveal a main effect of *Stimulation Type*, suggesting that active rTMS did not improve accuracy over sensory-matched electrical sham stimulation. Nevertheless, the results revealed an overall pattern that was consistent with the a priori expectations that rTMS has the largest modulatory effect for the hardest difficulty level of the hardest task condition (Invalid). Consequently, even if not significant, this improvement was considered in isolation to address a number of potentially relevant methodological considerations when applying rTMS for cognitive benefits, including difficulty titration, fMRI activations, and rTMS targeting based on fMRI activations and electric-field modeling. These parameters are reviewed further below.

The present design was based on a targeting approach in which parametric fMRI activations associated with increased set sizes were used to define the target of stimulation. Subsequent analyses of the accuracy data revealed this parametric fMRI activity negatively correlated with the improvement induced by active rTMS in the hardest difficulty level of the Invalid condition. An increase in the BOLD response is often associated with an increase in cognitive “effort” (Engström et al., 2013) and, conversely, a decrease in the BOLD signal is often interpreted as an increase in “efficiency” of the area in performing its function. This result supports the state-dependency assumption defined by Silvanto and collaborators (Silvanto et al., 2008) stating that rTMS is more effective for the most deactivated neural populations, and extends it to higher cognitive tasks than the perceptual tasks used in their studies.

The target of stimulation was selected according to individual brain activations and refined using electric field modeling to elicit the strongest magnetic field on the stronger brain activation obtained while subjects were performing the task. While this approach is more sophisticated than conventional rTMS targeting, it is unclear whether the rTMS effects were enhanced as a result, since it was not feasible to implement a comparison targeting condition in this study. However, by plotting the individual coil location and orientation for each individual subject, we were able to observe that this targeting method led to large spread in area of stimulation, which could have diluted the effects of rTMS even though the subsequent analysis did not reveal any correlation between rTMS effect and individual Euclidian distance to the group (using fMRI activation and coil location). If the individualized targeting method had been proven to be more effective to induce stronger rTMS effects (Sack et al., 2009), it is possible that the method chosen in this study was not the most effective, and a new alternative needs to be developed to better account for between-subjects’ variability.

Finally, it is worth nothing that the stimulation frequency (5Hz) and intensity of stimulation (100% rMT) were selected as they have previously been shown to be effective for inducing cognitive enhancement (Luber et al., 2007). Individualizing these parameters by using closed-loop information from simultaneously-recorded EEG may be a potential way to achieve greater specificity and achieve a larger effect size. Future studies may wish to attempt this approach. Lastly, previous TMS work on WM maintenance has shown improvements primarily with posterior targets such as the parietal cortex (Hamidi et al., 2008; Luber et al., 2007; 2008) making these also putative targets for future attempts to enhance WM manipulation.

## Conclusions

Effect of active and sham 5Hz rTMS on accuracy interacted with task difficulty in a working memory manipulation task, leading to a slight performance improvement in the hardest condition. The magnitude of this improvement was moderated by baseline activity within the DLPFC, while the timing of stimulation relative to encoding and retrieval in the task did not influence the rTMS effects. The effects of online rTMS should be investigated in older populations under highly challenging memory demands as it could be a promising method to reduce the cognitive decline associated with healthy aging and neurodegenerative disorders.

## Acknowledgments

The authors would like to thank Joyce Wang, Garland Austin, Laura Holton, and Annie Apple for help with piloting of preliminary participants; Alexandra Brito and Hannah Palmer for help with manuscript preparation; Emelina Vienneau for assistance in setting up the E-field modeling pipeline; Stefan M. Goetz for help with the robot configuration; and Alaattin Erkanli for help with data analysis.

